# IL-17RA-signaling modulates CD8+ T cell survival and exhaustion during *Trypanosoma cruzi* infection

**DOI:** 10.1101/314336

**Authors:** Jimena Tosello Boari, Cintia L. Araujo Furlan, Facundo Fiocca Vernengo, Constanza Rodriguez, María C. Ramello, María C. Amezcua Vesely, Melisa Gorosito Serrán, Nicolás G. Nuñez, Wilfrid Richer, Eliane Piaggio, Carolina L. Montes, Adriana Gruppi, Eva V. Acosta Rodríguez

## Abstract

The IL-17 family contributes to host defense against many intracellular pathogens by mechanisms not fully understood. CD8+ T lymphocytes are key elements against intracellular microbes and their survival and appropriate response is orchestrated by several cytokines. Here, we demonstrated that IL-17RA-signaling cytokines sustain pathogen-specific CD8+ T cell immunity. Absence of IL-17RA and IL-17A/F during *Trypanosoma cruzi* infection resulted in increased tissue parasitism and reduced frequency of parasite-specific CD8+ T cells. Impaired IL-17RA-signaling *in vivo* increased apoptosis of parasite-specific CD8+ T cells while recombinant IL-17 *in vitro* down-regulated the pro-apoptotic protein BAD and promoted activated CD8+ T cell survival. Phenotypic, functional and trancriptomic profiling showed that *T. cruzi*-specific CD8+ T cells arising in IL-17RA-deficient mice presented features of cell dysfunction. PD-L1 blockade partially restored the magnitude of CD8+ T cell responses and parasite control in these mice. Adoptive transfer experiments established that IL-17RA-signaling is intrinsically required for the proper maintenance of functional effector CD8+ T cells. Altogether, our results identify IL-17RA and IL-17A as critical factors for sustaining CD8+ T cell immunity to *T. cruzi*.

## Introduction

IL-17A and IL-17F were initially associated to the pathogenesis of autoinflammatory and autoimmune disorders [1]. Nonetheless, the primary function of these cytokines is likely host protection against microbes. Indeed, IL-17A and/or IL-17F-deficient mice are highly susceptible to a wide array of infections with fungi and extracellular bacteria but also with viruses and parasites [2,3]. We previously demonstrated that cytokines of the IL-17 family play a critical role in host survival by regulating exuberant inflammation and immunopathology during infection with the protozoan *Trypanosoma cruzi* [4,5]. Additional data from our laboratory suggested that the IL-17 cytokines played additional protective roles against *T. cruzi* by modulating adaptive immunity.

The IL-17 family is constituted by 6 members: IL-17A to IL-17F that have different cellular sources and expression patterns but often show overlapping activities. They signal through a receptor complex composed of at least two identical or different subunits of the IL-17R family (IL-17RA to IL-17RE). IL-17RA is the common signaling subunit used by at least four ligands: IL-17A, IL-17C, IL-17E and IL-17F. IL-17 members regulate inflammation by recruiting and activating neutrophils, NK cells and other cells of the innate immune system and by inducing several pro-inflammatory mediators (i.e. cytokines, chemokines, microbial peptides and metalloproteinases). In addition, these cytokines play important roles during adaptive immune responses including the modulation of germinal center reactions as well as the regulation of Th1 and CD8+ cellular responses [6-9]. Infection with *T. cruzi* causes Chagas’ disease that is the third most frequent parasitic disease worldwide, endemic in Latin America and increasing globally due to migratory flows. Disease progression, from symptomless to severe cardiac and digestive forms, is linked to parasite heterogeneity and variable host immune response. CD8+ T cell mediated immunity is essential for parasite control throughout all the stages of the infection, although not sufficient for complete parasite elimination [10,11]. CD8+ T cell deficient mice are extremely susceptible to infection [12,13] and strategies that improve specific CD8+ T cell responses result in increased host protection [11,14,15]. Considering this and that parasite persistence correlates with disease severity in experimental and human *T. cruzi* infection [16,17], many efforts are focused at understanding the mechanisms that promote protective anti-parasite CD8+ T cell immunity.

The general features of protective CD8+ T cell responses (as defined with model viral and bacterial infections) consist of the generation and expansion of short-lived, highly functional effector populations that operate to clear the microbes. After pathogen control, most of the effector cells die and the few remaining cells differentiate into memory T cells, contributing to long-lived immunological protection [18]. During certain chronic infections, persistently activated CD8+ T cells acquire a state of cell dysfunction characterized by the hierarchical loss of effector functions and proliferation potential accompanied by sustained expression of multiple inhibitory receptors. Severe exhaustion leads to pathogen-specific cells with reduced ability to produce cytokines or to degranulate and prone to deletion. These exhausted CD8+ T cells fail to provide optimal protection contributing to poor pathogen control [19]. Antigen-specific and costimulatory signals are the driving forces of CD8+ T cell responses, but also cytokines (i.e. IL-21, IL-2, IL-12) are required to provide the “third-signal” essential for protective CD8+ T cell immunity [20,21].

Herein, we demonstrate that IL-17RA signaling is required for the maintenance of robust specific CD8+ T cell responses and host resistance to *T. cruzi*. We show that absence of IL-17RA signaling during this infection alters the transcriptional program of the effector CD8+ T cells affecting their survival, effector function and exhaustion. Our findings are of relevance for a better understanding of the role of IL-17 family on the orchestration of protective immunity against infections.

## Results

### IL-17RA and IL-17A/IL-17F deficiencies compromise parasite control and reduce the magnitude of the specific CD8+ T cell response during *T. cruzi* infection

We previously showed that IL-17RA-signaling and IL-17-secreting B cells are critically required for host survival during *T. cruzi* infection by, at least in part, regulating innate immunity and inflammation [4,5]. In addition, histological data from these initial studies suggested that infected IL-17RA knockout (KO) mice exhibited increased parasite levels in tissues but similar parasitemia than infected wild-type (WT) counterparts [4]. To confirm this and study possible additional protective mechanisms mediated by IL-17, we evaluated tissue parasitism in infected wild-type (WT) and IL-17RA knockout (KO) mice. As determined by the amounts of parasite DNA quantified in spleen, we observed that *T. cruzi* infected IL-17RA KO mice controlled tissue parasitism similar to WT mice during the first two weeks of infection (Fig 1A). However, IL-17RA KO mice failed to reduce parasite burden at later stages of infection, as attested by the increased parasitism in the spleen at 22 and 130 day post-infection (dpi) (Fig 1A). Furthermore, increased parasite levels were also detected in other organs such as liver and heart (Fig 1B-C).

**Fig 1.**
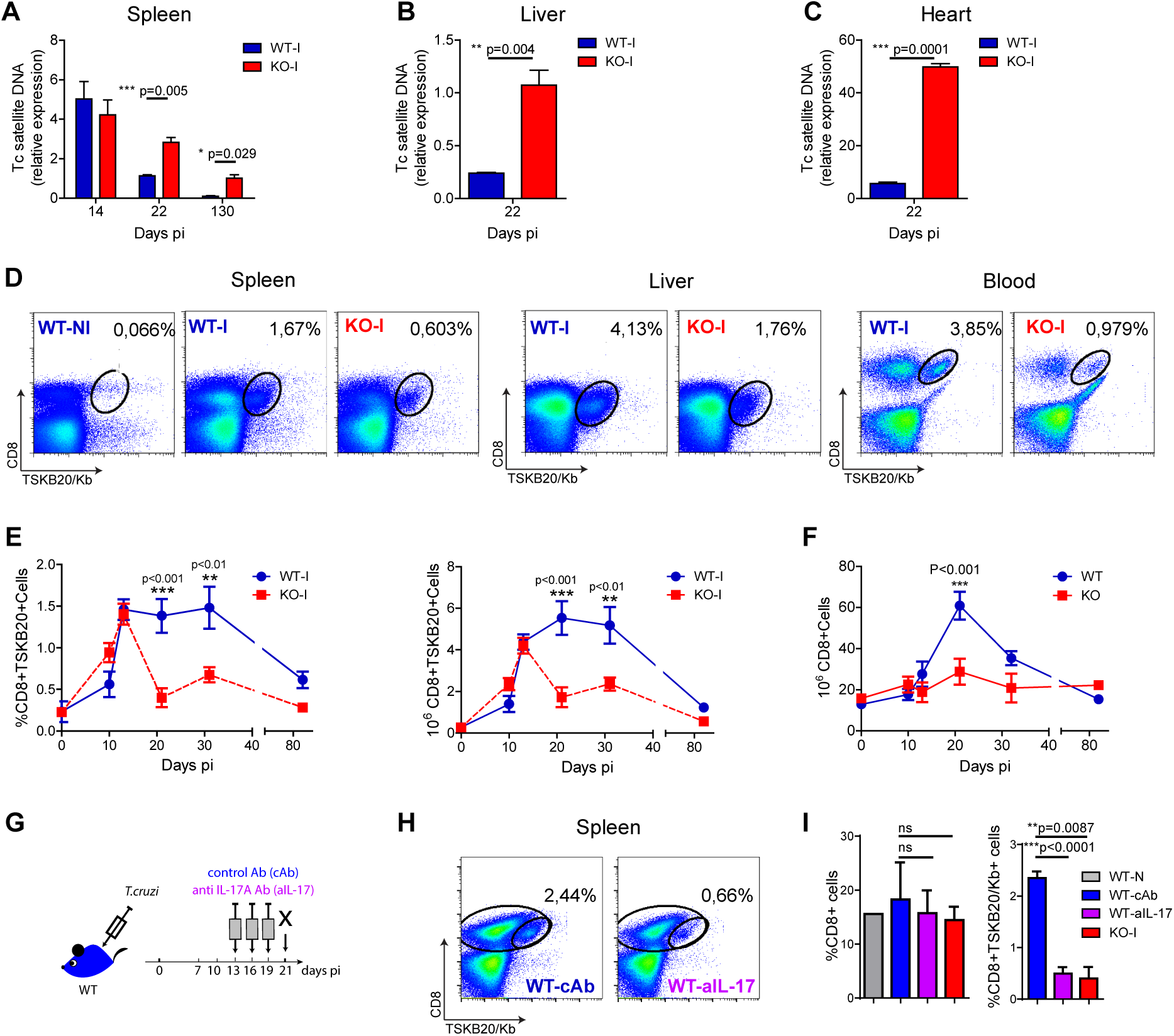
Absence of IL-17RA signaling results in increased tissue parasitism and a reduced magnitude of parasite-specific CD8+ T cell responses. (**A-C**) Relative amount of *T. cruzi* satellite DNA in spleen (**A**), liver (**B**) and heart (**C**) of infected WT and IL-17RA KO (KO) mice determined at the indicated dpi. Murine GAPDH was used for normalization. Data are presented as mean ± SD, N=5 mice. P values calculated with two-tailed T test. (**D**) Representative plots of CD8 and TSKB20/Kb staining in spleen, liver and blood of WT and KO mice at 22 dpi (WT-I and KO-I, respectively). A representative plot of the staining of splenocytes from non-infected WT mice (WT-N) is shown for comparison. (**E**) Percentage and cell numbers of TSKB20/Kb+ CD8+ T cells and (**F**) cell numbers of total CD8+ T cells determined in spleen of WT and IL-17RA KO mice at different dpi. Data shown as mean ± SD, N=5–8 mice. P values calculated using two-way ANOVA followed by Bonferroni‘s post-test. (**G**) Experimental layout of IL-17 neutralization in infected WT mice injected with control isotype or anti-IL-17 Abs (WT-cAb and WT-aIL-17, respectively). (**H**) Representative plots of CD8 and TSKB20/Kb staining in spleen and liver infected WT treated as indicated in G. (**I**) Percentage of total and TSKB20/Kb+ CD8+ T cells in mice treated as depicted in G. Results from non-infected WT mice (WT-N) and infection-matched KO mice (KO-I) are shown for comparison. Data are representative of five (**A-F**), and two (**G-I**) independent experiments.

As the aforementioned result suggested a deteriorated adaptive immune response, we focused at studying the role of IL-17RA in the development of parasite-specific CD8+ T cells. To this end, we examined the generation of CD8+ T cells specific for the immunodominant epitope TSKB20 *(T. cruzi* trans-sialidase amino acids 569-576 – ANYKFTLV–) [22] in infected IL-17RA KO and WT mice. We determined that the frequency of TSKB20-specific CD8+ T cells was reduced in spleen, liver and blood of infected IL-17RA KO mice at 20 dpi (Fig 1D). Kinetics studies showed that the frequency and absolute numbers of spleen TSKB20-specific CD8+ T cells detected in infected IL-17RA KO mice was similar to that infected WT controls up to 14 dpi, but these values dropped dramatically soon after (Fig 1E). Remarkably, total CD8+ T cell numbers at 20 dpi were also reduced in the spleen of infected IL-17RA KO mice (Figure 1F). In agreement with the notion that IL-17A and IL-17F are the major IL-17RA-signaling cytokines able to modulate immune responses, IL-17A/IL-17F double knockout (DKO) mice also showed increased tissue parasitism in spleen and liver (Fig S1A) and reduced frequency of parasite-specific CD8+ T cells in comparison to WT controls (Fig S1B).

Considering the kinetics of the TSKB20-specific CD8+ T cell immunity observed in infected IL-17RA KO mice (Fig 1E) as well as our reported results showing that IL-17 production peaked around 14 dpi in this experimental infection model [4,5], we speculated that IL-17RA signaling plays a role after priming, during the expansion and maintenance of the parasite-specific CD8+ T cells. To test this, we performed neutralization experiments in which infected WT mice were injected with anti-IL-17A Abs from 13 dpi and during a limited period (Fig 1G). Blockade of IL-17 signaling specifically during the expansion/maintenance phase resulted in a significant decrease in the frequency of TSKB20-specific T cells (Fig 1H). Indeed, mice treated with anti-IL-17 only after the CD8+ T cell response was already established (from 13 to 19 dpi), showed a frequency of parasite-specific T cells comparable to that observed in infected IL-17RA KO mice that lacked IL-17 signaling throughout the infection (Fig 1I).

### Absence of IL-17RA signaling during *T. cruzi* infection reduces survival of total and specific CD8+ T cells

Considering the conventional kinetics of T cell responses [18], it was conceivable that the abortive CD8+ T cell response observed in infected IL-17RA KO mice could be a consequence of a reduced cell expansion and/or increased contraction. To address the role of IL-17RA in CD8+ T cell proliferation we performed *in vivo* experiments of BrDU incorporation between days 15 to 20 dpi, time points that directly preceded the dramatic decrease in the numbers of specific CD8+ T cells in infected IL-17RA KO mice. Results depicted in Fig 2A demonstrated that absence of IL-17RA does not compromise proliferation of CD8+ T cells during *T. cruzi* infection. Complementary studies of ex *vivo* detection of the Ki-67 antigen led to similar conclusions (Fig S2A). Then, we tested a possible role of IL-17RA signaling in supporting CD8+ T cell survival. In comparison to counterparts from infected WT mice, total and TSKB20-specific CD8+ T cells from infected IL-17RA KO mice showed significantly higher frequencies of apoptotic cells identified by TMRE-based mitochondrial membrane potential assay as early as 10 dpi and also at later time points (20 dpi) (Fig 2B). Similar results were obtained by analyzing Annexin V+/7AAD-cells (Fig S2B). In the same direction, total and parasite-specific CD8+ T cells from infected WT mice treated with anti-IL-17A Abs (Fig 1H) showed slightly higher percentage of apoptotic cells in comparison to counterparts from controls \, reaching levels of apoptosis similar to that determined in CD8+ T cells from infected IL-17RA KO mice (Fig S2C).

**Fig 2.**
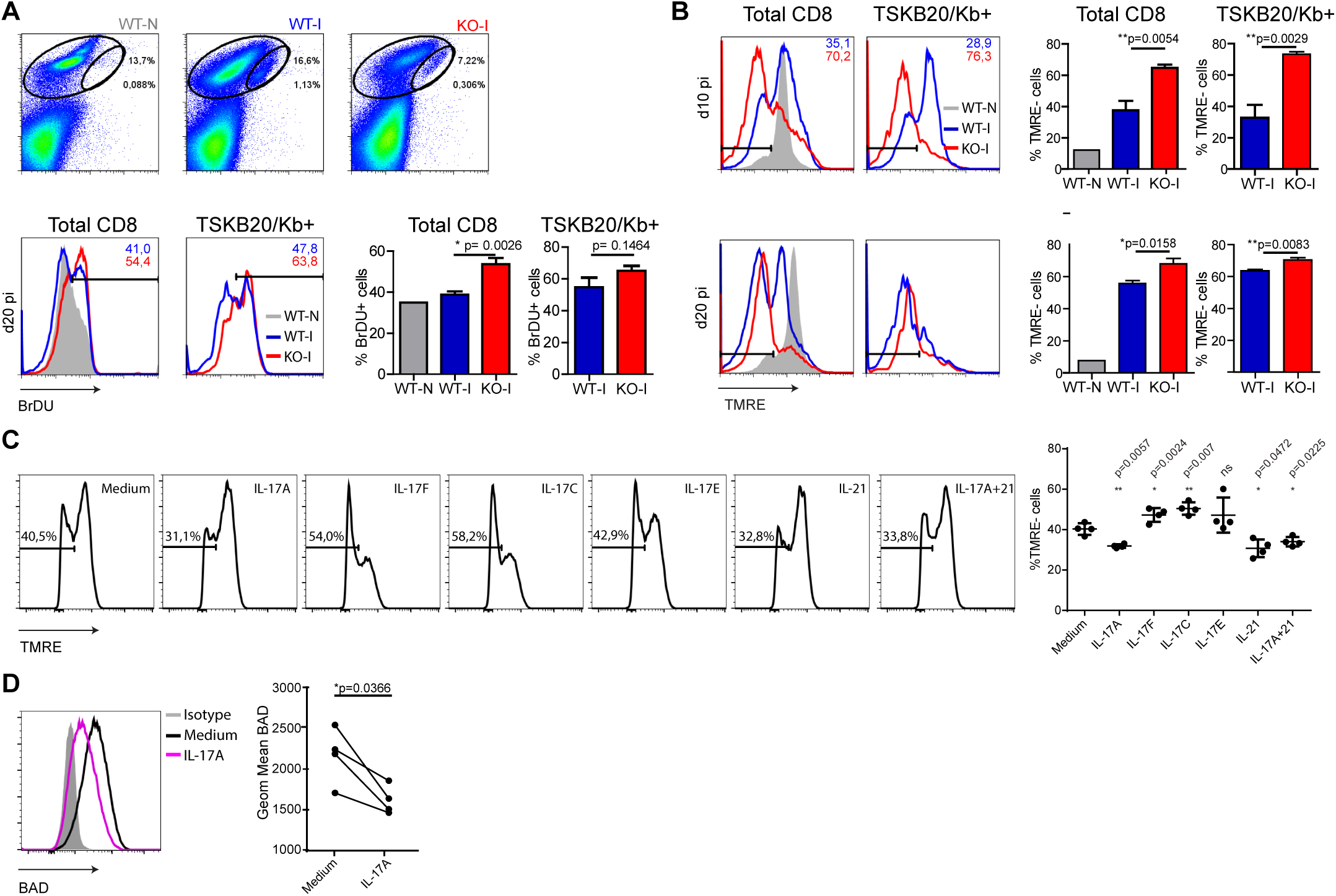
IL-17RA signaling during *T. cruzi* infection promotes survival of CD8+ T cells. (**A**) Representative dot plots of CD8 and TSKB20/Kb staining in spleen of WT and KO mice at 0 dpi (WT-N) and 20 dpi (WT-I and KO-I) and representative histograms of BrDU staining in the indicated total and TSKB20/Kb+ CD8+ T cell gate. Numbers indicate the frequency of BrDU+ cells from the corresponding colored group. Histograms are representative of one out of five mice. Bars in the statistical analysis represent data as mean ± SD, N=5 mice. P values were calculated with two-tailed T test. (**B**) Representative histograms of TMRE staining in total (left) and TSKB20/Kb+ (right) CD8+ T cells from the spleen of WT and IL17RA KO mice at 10 dpi (top) and 20 dpi (bottom). Numbers indicate the frequency of TMRE- (apoptotic) cells in total and TSKB20/Kb+ CD8+ T cells from the corresponding colored group. Grey tinted histogram show staining in CD8+ T cells from non-infected WT mice (WT-N). Histograms are representative of one out of five mice. Bars in the statistical analysis represent data as mean ± SD, N=5 mice. P values were calculated with two-tailed T test. (**C** and **D**) Representative histograms of TMRE (**C**) and BAD (**D**) stainings in cultures of purified CD8+ T cells activated with coated anti-CD3 and anti-CD28 in the presence of the indicated cytokines. Statistical analysis in (**C**) shows the frequency of TMRE- (apoptotic) cells in each biological replicate (N=4 mice) and the mean ± SD. P values calculated with paired two-tailed T test (ns: not significant). Statistical analysis in (**D**) represent protein expression as the geometric mean of fluorescence intensity in each replicate (N=4 mice). Lines link paired samples. P values calculated with paired two-tailed T test. Data in **A-C** and **D** are representative of three and two independent experiments, respectively.

Given the results described above, we evaluated whether cytokines that signal through IL-17RA were able to directly increase survival of activated CD8+ T cells. CD8+ T cells purified from the spleen of non-infected WT mice were activated during 24 h with plastic-coated anti-CD3 and anti-CD28 Abs in the presence of recombinant cytokines. Addition of IL-17A but not IL-17F, IL-17C and IL-17E lead to a modest but significant reduction in the percentage of apoptotic activated CD8+ T cells (Fig 2C). Of note, the direct anti-apoptotic effect of IL-17 in CD8+ T cells was comparable to that of IL-21, a well-recognized survival factor for this population [23]. Trying to elucidate the mechanisms underlying the pro-survival effects of IL-17A, we evaluated the expression of pro and anti-apoptotic members of the Bcl-2 superfamily that are critically involved in the regulation of T cell death [24]. We determined that IL-17A significantly downregulated the expression of Bad (Fig 2D). Very small or no changes were induced in other members such as Bcl-xL, Bcl-2, Bim and Bax (Fig S2D). Altogether, these results indicate that IL-17A may promote CD8+ T cell survival by down-regulating pro-apoptotic factors of the Bcl-2 family.

### Absence of IL-17RA perturbs the transcriptional program of CD8+ T cells activated during *T. cruzi* infection

To gain further insights of the overall impact of the absence of IL-17RA signaling, we determined the gene expression profiles of purified CD8+ T cells using Affymetrix microarrays. By normalizing gene expression to the corresponding non-infected samples, we compared the transcriptional changes induced by the infection in CD8+ T cells from IL-17RA KO mice versus counterparts from WT mice in a fold change/fold change plot. We identified 4287 genes that showed a statistically significant difference in expression of over 1.2 fold (p<0.05) and were grouped into different sets according to the differential pattern of expression (Fig 3A). Remarkably, 1647 genes (sets pink, orange and red) were expressed at higher level while 2640 genes (sets light green, green and dark) were expressed at lower levels in CD8+ T cells from IL-17RA KO infected mice in comparison to WT counterparts. To further evaluate the biologic characteristics of the transcriptional landscape of CD8+ T cells from infected IL-17RA KO and WT mice, we performed gene-enrichment set analysis (GSEA) in all the genes differentially expressed. After reaching broad results using gene ontology, immunological and curated databases, we decided to focus on specific gene sets (Molecular Signatures DB-MSigDB) related with CD8+ T cell function and differentiation such as GSEA26495 and GSEA41867. This supervised analysis showed that IL-17RA KO CD8+ T cells were significantly enriched in six gene sets within which the most significant enriched CD8+ T cell gene sets in the significance order (size of FWER P values) were PD1^hi^ versus PD1^low^ (p<0.001) and Naïve versus PD1^low^ (p<0.001) (Fig 3B). These results denoted that CD8+ T cells elicited during *T. cruzi* infection in absence of IL-17RA showed a phenotype with mixed features of naïve and dysfunctional cells.

**Fig 3.**
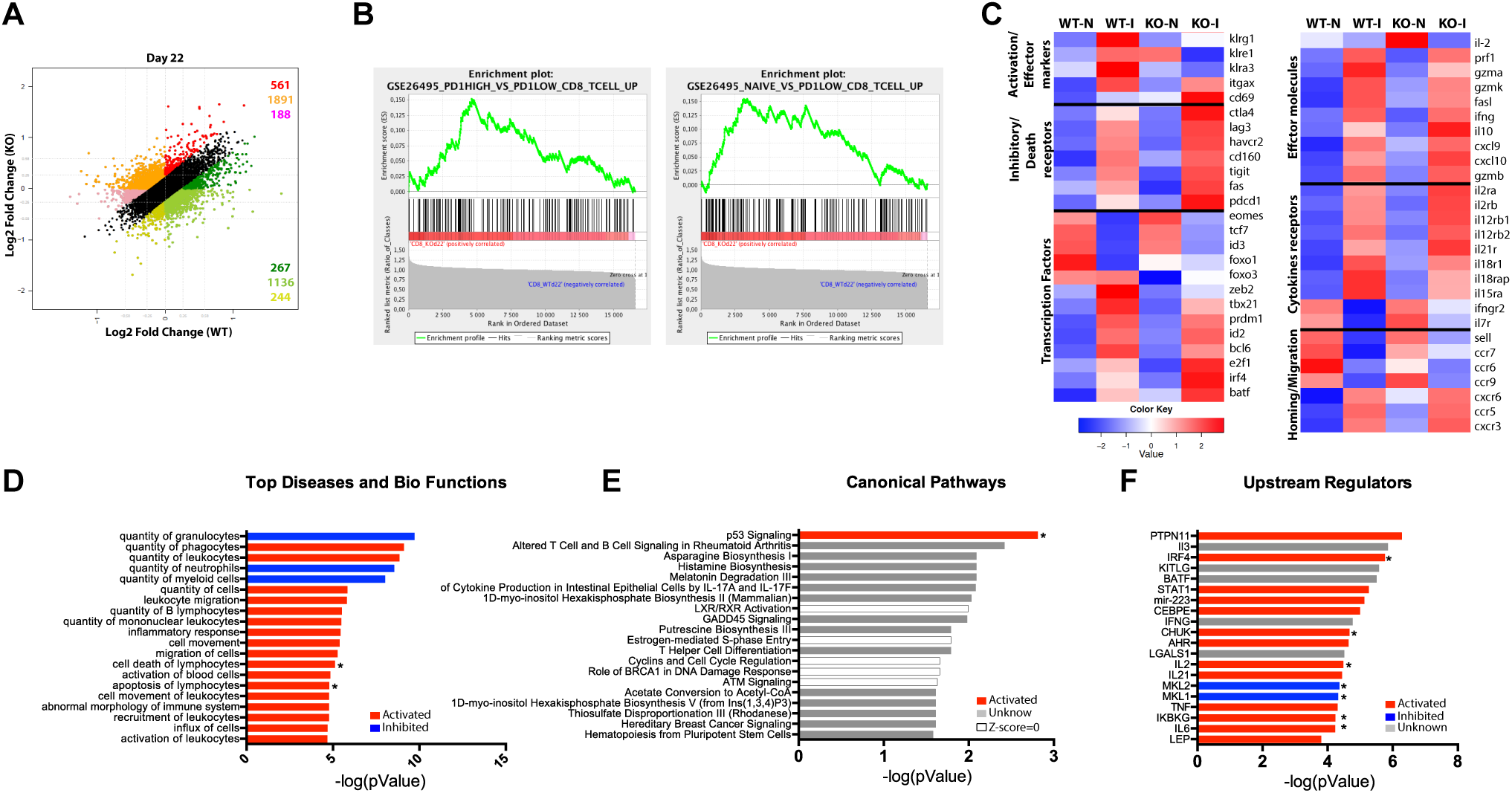
Substantial differences in the gene-expression profile of CD8+ T cells from *T. cruzi*-infected IL-17RA KO and WT mice. (**A-F**) Microarray analysis of purified CD8+ T cells from infected WT (WT-I) and IL-17RA KO (KO-I) mice (22dpi) and non-infected counterparts (WT-N and KO-N). N=3 mice per group. (**A**) Dot plots displaying the number of genes that show a significant ≥1.2-fold difference in fold change expression of WT-I and KO-I relative to non-infected counterparts. Colors indicate sets of genes with different expression patterns. (**B**) Top two enrichment plots in KO-I (p<0.001) determined by supervised analysis of all infection induced genes in WT-I and KO-I using GSEA26495 and GSEA41867 (MSigDB C5BP) signature gene sets. (**C**) Heat maps of expression of selected genes according to categories relevant to CD8+ T cell biology. Each gene expression value was represented by the median value from 3 mice. (**D-F**) IPA of genes induced by the infection but showing significantly higher expression in IL-17RA KO mice (red genes in A). Top 20 “Diseases and Bio-functions” (**D**), “Canonical pathways” (**E**) and “Up-stream regulators” (**F**) that were most significantly altered (P-value < 0.05 with Fischer’s exact test) in KO mice are shown. The activation Z-score was calculated to predict activation (red), inhibition (blue), categories where no predictions can be made and unknown results (with Z-score close to 0). Categories significantly activated or inhibited according to Z-score are marked with a star.

Given the previous results, we then analyzed the expression patterns of genes selected according to categories relevant for CD8+ T cell biology. As observed in the heat maps, gene expression profiles of CD8+ T cells from WT and IL-17RA KO mice were similar before infection but changed significantly after 22 dpi (Fig 3C). Indeed, genes encoding activation and effector cell markers such as the KLRG family molecules, CD11c and CD69 showed completely reciprocal pattern of expression in CD8+ T cells from infected IL-17RA KO versus WT mice. Furthermore, CD8+ T cells from infected IL-17RA KO mice showed clearly increased levels of many genes encoding inhibitory and death receptors including *Ctla4, Lag3, Tim3, Pd1, Cd160, Tigit*, and *Fas*. We also analyzed how IL-17RA signaling modulated the expression of genes encoding TFs that regulate CD8+ T cell fate. Of note, there was a group of TF-encoding genes that were significantly more up-regulated in CD8+ T cells from infected IL-17RA KO mice. This group included genes such as *Irf4* and *Batf* that are critical for the generation of effector CD8+ T cells [25-27]. Finally, we determined that CD8+ T cells from infected WT and IL-17RA mice showed differences in the expression of many genes encoding effector molecules, cytokine receptors and homing and migration molecules that in overall supported the fact that absence of IL-17RA influence CD8+ T cell fate.

To broadly analyze the significance of the transcriptional differences between CD8+ T cells from infected IL-17RA KO mice versus WT counterparts, we performed an Ingenuity Pathway Analysis (IPA) on genes marked in red in Figure 3A, which were up-regulated by the infection in the CD8+ T cells from both groups of mice but were significantly expressed at higher levels in CD8+ T cells from IL-17RA KO mice. The top 20 “Diseases and Bio-functions”, “Canonical pathways” and “Up-stream regulators” that were most significantly altered in CD8+ T cells elicited by *T. cruzi* infection in the absence of IL-17RA are shown in the Fig 3D. Remarkably, “Cell death” and “Apoptosis of Lymphocytes” bio-functions and the “p53 Signaling” canonical pathway emerged as significantly up-regulated. These results support the notion that lack of IL-17RA signaling during *T. cruzi* infection significantly affected the survival/senescence status and also participated in several biological features of the CD8+ T cells. Finally, to elucidate which molecule/s could govern the observed differences in gene expression, we performed an “upstream regulator analysis”. Many modulators of the inflammatory responses including the cytokines IL-2 and IL-6 and the regulators of NFkB as CHUK and IKBKG were significantly up-regulated while MKL-2 and MKL-1 that interacts with transcriptional regulator serum response factor were down-regulated in CD8+ T cells from infected IL-17RA KO mice. Remarkably, IRF4 and at a minor extent other TF such as BATF, STAT1 and AHR emerged as potential upstream regulators of the transcriptional program of the CD8+ T cells induced by *T. cruzi* infection in absence of IL-17RA signaling.

### Absence of IL-17RA signaling during *T. cruzi* infection results in CD8+ T cell exhaustion

Given microarray data and considering that the increased T cell deletion could be consequence of a process of cell exhaustion, we analyzed the expression of inhibitory and death receptors in CD8+ T cells from infected WT and IL-17RA KO mice. Compared to CD8+ T cells from non-infected mice, total and TSKB20-specific CD8+ T cells from infected WT mice presented a slight upregulation of PD-1 and TIGIT as well as similar frequency of CTLA-4+ cells and reduced frequency of BTLA+ cells (Fig 4A). Noteworthy, total and, particularly, TSKB20-specific CD8+ T cells from infected IL-17RA KO mice showed significantly higher expression of PD-1 and TIGIT together with increased frequency of CTLA-4+ and BTLA+ cells. Furthermore, these cells presented an apoptosis-prone phenotype characterized by the expression of high levels of CD95/Fas and CD120b/TNF-R2 (Figure 4B). Total and TSKB20-specific CD8+ T cells from infected IL-17A/IL-17F DKO mice showed a profile of expression of inhibitory (PD-1) and death receptors (Fas/CD95) that was comparable to that of counterparts from infected IL-17RA KO mice (Fig S1C and D).

**Fig 4.**
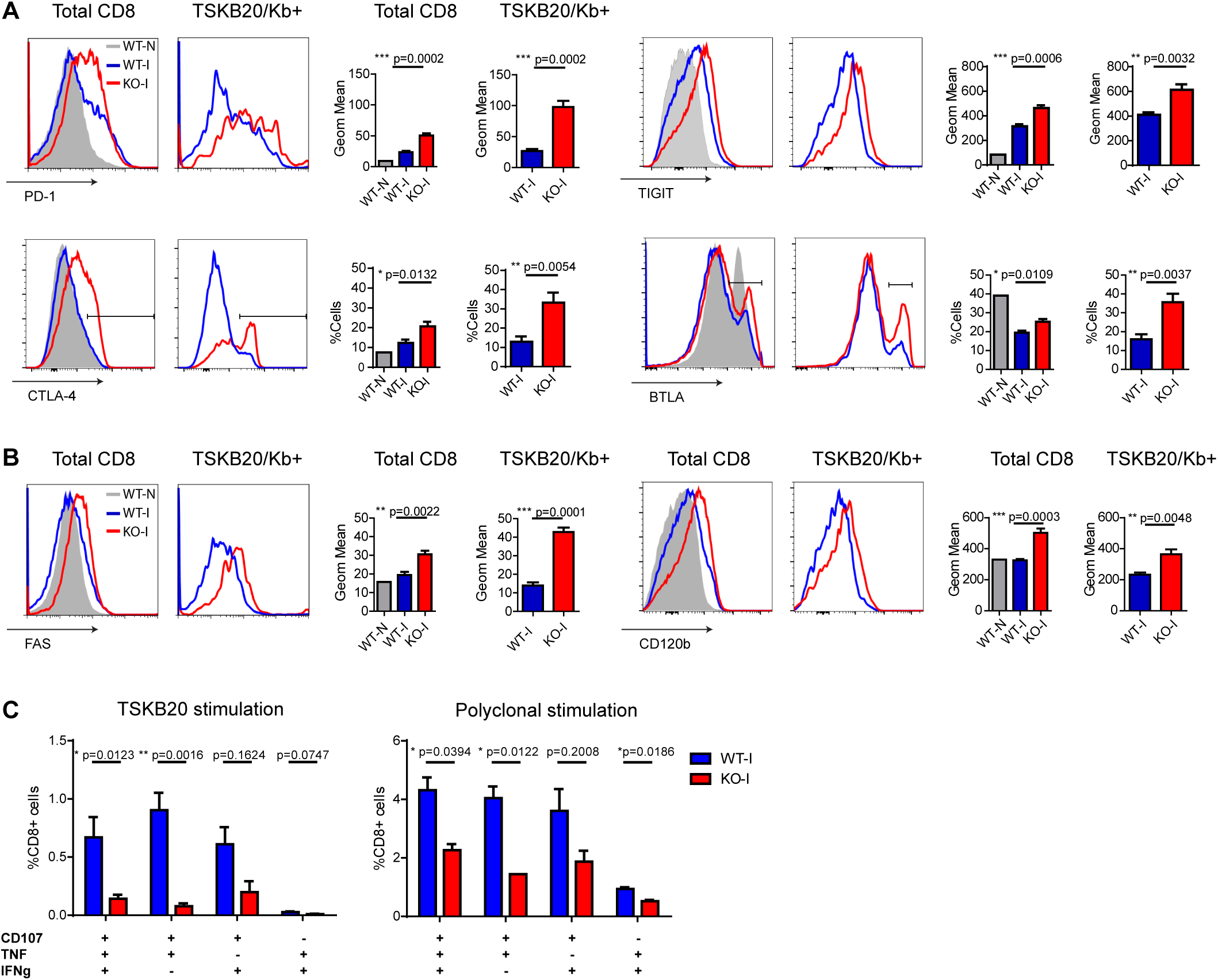
CD8+ T cells from *T. cruzi* infected IL-17RA KO mice show features of exhausted cells. (**A-B**) Representative histograms and statistical analysis of the geometric mean of expression or percentage of cells expressing several inhibitory **(A)** and death receptors **(B)** in total and TSKB20/Kb+ spleen CD8+ T cells from infected WT (WT-I) and IL-17RA (KO-I) mice (22 dpi). Staining of non-infected WT mice (WT-N) is showed as gray tinted histograms for comparison. (**A-B**) Data in statistical analysis are presented as mean ± SD, N=4–6 mice. **(C)** Percentage of spleen CD8+ T cells from infected WT and KO mice (22 dpi) that exhibit polyfunctional effector function denoted by expression of CD107a, IFNγ and/or TNF upon 5 h of the indicated stimulation. Data shown as mean ± SD, N=5 mice. Data were background subtracted. All P values were calculated using two-tailed T test. Data are representative of at least three independent experiments.

To address whether the phenotypic features of CD8+ T cells elicited in absence of IL-17RA correlated with altered functionality, we compared the effector function of CD8+ T cells from infected WT and IL-17RA KO mice. To this end, we analyzed the mobilization of CD107a and secretion of effector cytokines upon specific and polyclonal activation *in vitro*. Stimulation of splenocytes from infected IL-17RA KO mice with the TSKB20 peptide resulted in significant reduced frequency of CD8+ T cells showing polyfunctional characteristics (surface CD107a expression and IFNγ and/or TNF production) in comparison to WT controls (Fig 4C, left panel and Fig S3). Although a poor response was anticipated due to the low frequency of TSKB20-specific CD8+ T cells in infected IL-17RA KO mice, the antigen-specific effector response of CD8+ T cells was barely different from background. Stronger polyclonal stimulation with PMA and Ionomycin increased the percentage of CD8+ T cells within splenocytes from infected IL-17RA KO mice showing a polyfunctional response; however, the magnitude of this effector response was significantly lower than that of WT counterparts (Fig 4C, right panel and Fig S3).

To determine if the higher parasite burden in infected IL-17RA KO mice may underlie the upregulation of the inhibitory receptors observed in their CD8+ T cells, we evaluated the phenotype of CD8+ T cells from WT mice infected with increasing parasite doses. Remarkably, increasing loads of *T. cruzi* did not diminish the frequency of parasite-specific CD8+ T cells nor promoted the upregulation of the inhibitory and death receptors evaluated (Fig S4).

### Checkpoint blockade partially restores parasite-specific CD8+ T cell immunity and enhances parasite control in infected IL-17RA KO mice

CD8+ T cell exhaustion has been associated with poor microbial control during many infections as well as with tumor progression in cancer. In these settings, checkpoint blockade has shown success to restore the magnitude and functionality of effector T cells and to enhance the control of the insult [28]. Furthermore, PD-L1 blockade has been effective to prevent T cell apoptosis in murine and human sepsis [29,30]. Our results showing that lack of IL-17RA signaling during *T. cruzi* infection lead to poor CD8+ T cell effector function and parasite persistence, prompted us to evaluate whether checkpoint blockade could restore a resistant phenotype in infected IL-17RA KO mice. As PD-1 is highly expressed on parasite-specific CD8+ T cells from infected IL-17RA KO mice, we targeted this checkpoint receptor by injecting blocking anti PD-L1 Abs at 15 and 18 dpi, before the contraction of parasite-specific CD8+ T cell response (Fig 5A). We determined that PD-L1 blockade in infected IL-17RA KO significantly augmented the magnitude of the TSKB20-specific CD8+ T cell response by 21dpi, particularly in spleen and liver (Fig 5B). Similar response to treatment was observed in infected WT mice. Remarkably, the enhanced *T. cruzi*-specific CD8+ T cell immunity correlated with a clear reduction in the levels of tissue parasitism (Figure 5C). In addition, inhibition of PD-L1 also lead to a milder infection as highlighted by the significantly reduced levels of several damage biomarkers such as the transaminase aspartate aminotransferase (AST), and creatinin kinase total (CK) and myocardial band (CK-MB) (Fig 5D). The reduction in the severity of the infection (parasite persistence and tissue damage) upon PD-L1 checkpoint blockade was more impressive in IL-17RA KO mice as they show an extremely high susceptibility to *T. cruzi* infection in absence of treatment.

**Fig 5.**
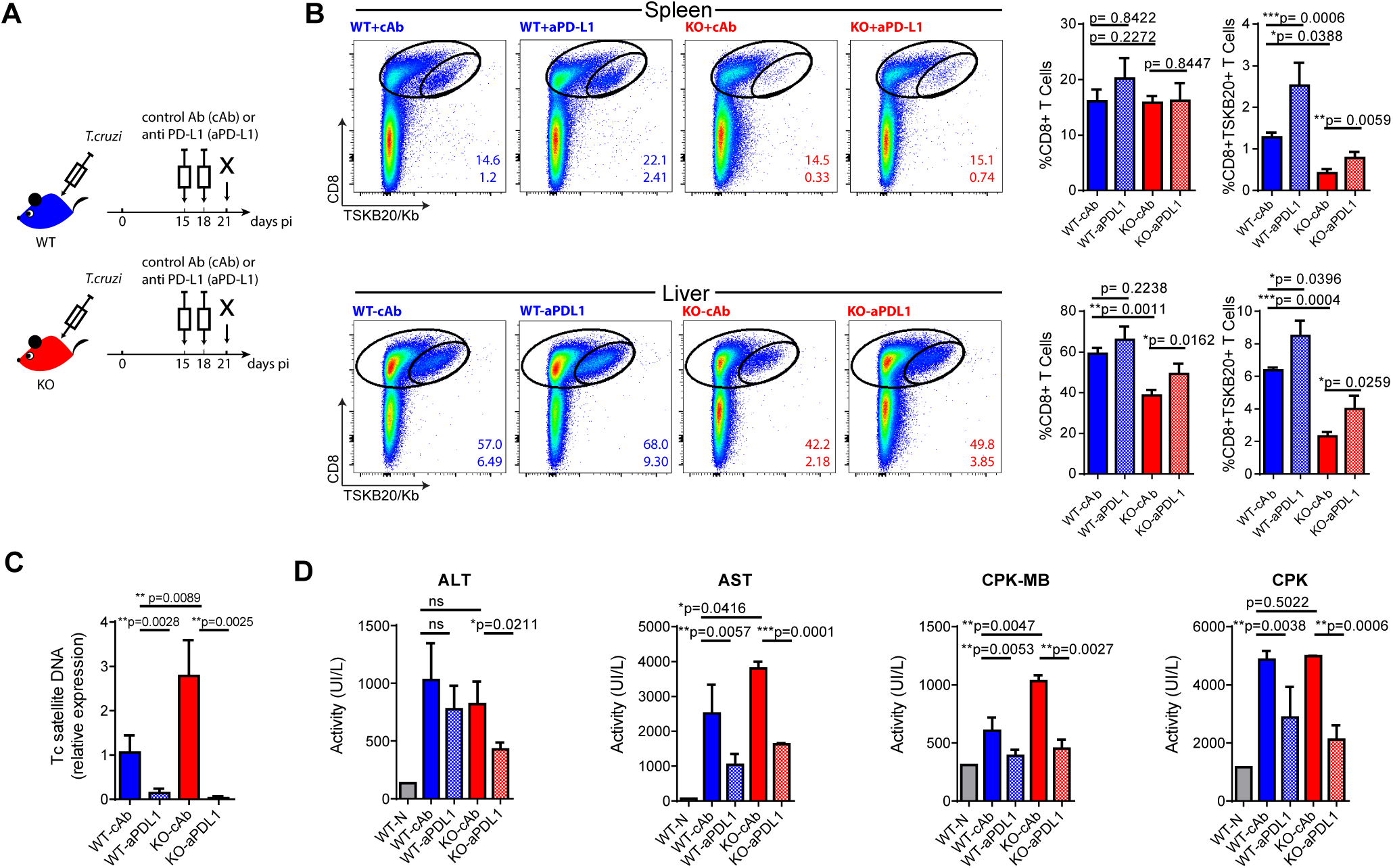
Checkpoint blockade reinvigorates parasite-specific CD8+ T cell immunity and enhances parasite control in infected IL-17RA KO mice. **(A)** Layout of the checkpoint blockade experiment. **(B)** Representative plots and statistical analysis of CD8 and TSKB20/Kb staining in spleen and liver of the experimental groups indicated in **(A)**. Numbers within plots indicate the frequency of total CD8+ cells (up) and TSKB20-specific CD8+ T cells (down) at 21 dpi. Bar graphs represent data as mean ± SD, N=7 mice. **(C)** Relative amount of *T. cruzi* satellite DNA in the spleens of the indicated experimental groups at 21 dpi. Murine GAPDH was used for normalization. Data are presented as mean ± SD relative to WT-cAb group, N=7 mice. **(D)** Enzymatic activity (UI/L) of alanine transaminase (ALT), aspartate transaminase (AST), creatinine kinase-MB and creatinine phosphokinase (CPK) on plasma of the experimental groups indicated in **(A)** at 21 dpi. N=7 mice. All P values were calculated with two-tailed T test. Data are representative of two independent experiments.

### Intrinsic IL-17RA signaling modulates the maintenance and phenotype of CD8+ T cells activated during *T. cruzi* infection

To understand whether IL-17RA played direct or indirect roles in the modulation of CD8+ T cell responses to *T. cruzi*, we first quantified in serum the concentration of IL-21, known to modulate CD8+ T cell immunity. Of note, we determined that in comparison to WT counterparts, infected IL-17RA KO mice exhibited increased concentration of IL-21 (Fig 6A). This result pointed out that high amounts of IL-21 are not able to compensate the lack of IL-17RA signaling for the induction of robust CD8+ T cell responses during this infection.

**Fig 6.**
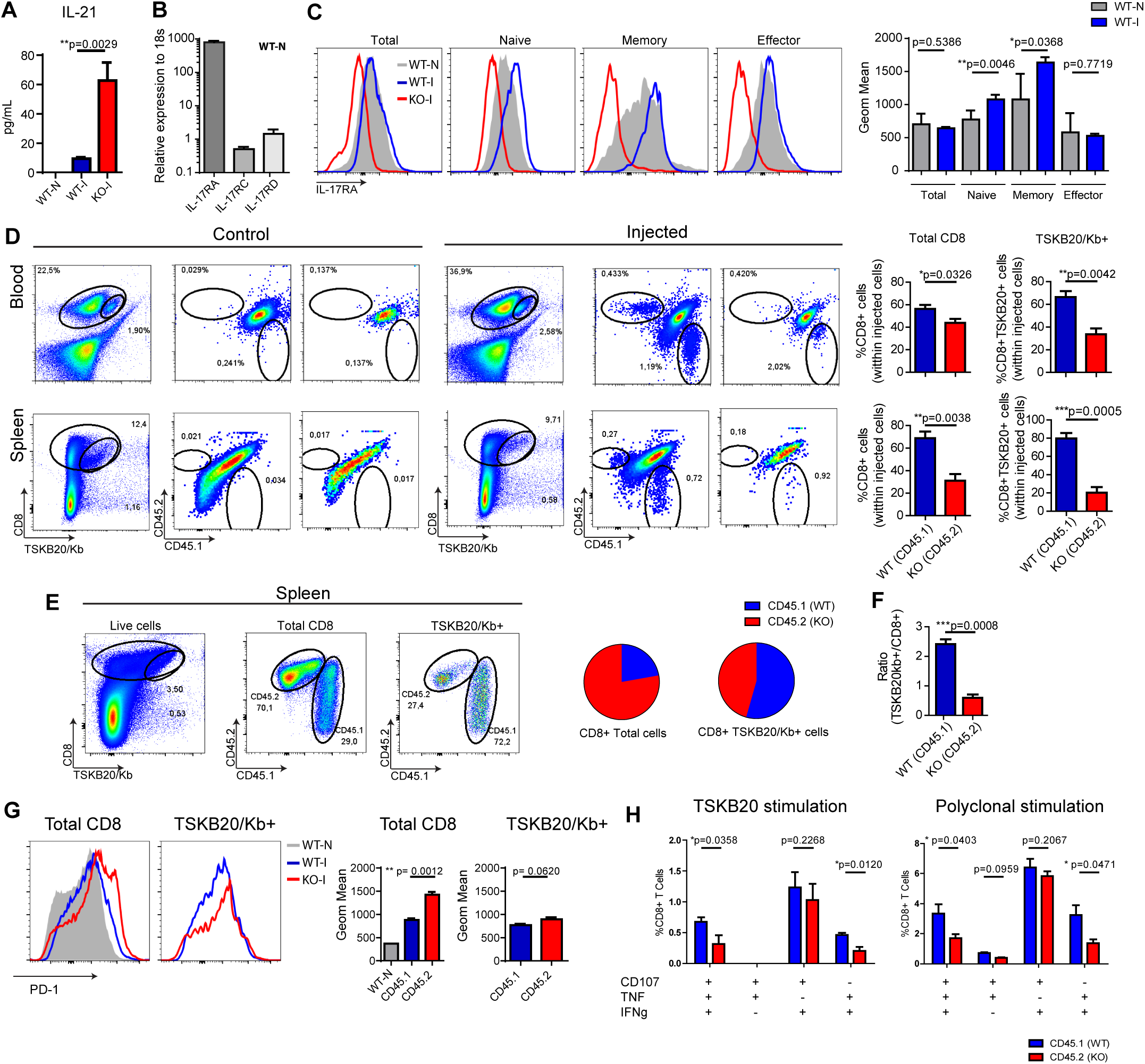
Intrinsic IL-17RA-signaling modulates the maintenance, phenotype and function of CD8+ T cells during *T. cruzi* infection. **(A)** Concentration of IL-21 in plasma of infected WT and IL-17RA KO mice at 20 dpi. Data are presented as mean ± SD, N=4 mice. **(B)** Amounts of *Il17ra, Il17rc* and *Il17rd* transcript determined in spleen CD8+ T cells purified from non-infected WT mice, normalized to 18S RNA. Data are presented as mean ± SD, N=4 mice **(C)** Representative histograms and statistical analysis of the expression of IL-17RA (protein) on different CD8+ T cell subsets identified according to CD44 and CD62L staining in spleen cell suspensions from non-infected **(N)** and infected **(I)** WT mice (22 dpi). Staining of IL-17RA on spleen CD8+ T cells from IL-17RA KO mice is showed as negative control. Data in statistical analysis are presented as mean ± SD, N≥6. **(D)** Representative plots and statistical analysis of CD8 and TSKB20/Kb staining and of CD45.1 and CD45.2 staining within Total and TSKB20/Kb+ CD8+ T cells in the blood and spleen of infected F1 CD45.1/CD45.2 WT recipient mice (20dpi) non-injected (control) or injected with equal numbers of CD45.1+ WT and CD45.2+ KO CD8+ T cells. Numbers in the plots indicate the frequency of the correspondent cell subset. Bar graphs display the frequency of CD45.1+ WT cells and CD45.2+ KO cells within the indicated populations upon gating only in the injected cells. **(E)** Representative plots and statistical analysis of CD8 and TSKB20/Kb staining and of CD45.1 and CD45.2 staining within Total and TSKB20/Kb+ CD8+ T cells in the spleen of infected CD8α-/- recipients (17dpi) injected with equal numbers of CD45.1+ WT and CD45.2+ KO CD8+ T cells. Pie charts display the frequency of CD45.1+ WT and CD45.2+ KO cells within the indicated gates. Bar graph shows the ratio between the frequencies of TSKB20/Kb+ CD8+ T cells and the total CD8+ T cells within CD45.1 WT and CD45.2 IL-17RA KO populations. **(F)** Representative histograms and statistical analysis of the geometric mean of PD-1 expression in total and TSKB20/Kb+ CD8+ T cells within CD45.1+ WT and CD45.2+ IL-17RA KO CD8+ T cells from the spleen of CD8α-/- recipient mice. Data in statistical analysis are presented as mean ± SD, N=4–6 mice. **(G)** Percentage of polyfunctional effector cells denoted by expression CD107a, IFNγ and/or TNF upon 5h of the indicated stimulation on CD45.1+ WT and CD45.2+ IL-17RA KO CD8+ T cells from the spleen of CD8α-/- recipient mice. Data shown as mean ± SD, N=4 mice. Data were background subtracted. All P values were calculated using two-tailed T test. Data are representative of two independent experiments.

We next evaluated the possibility of a direct effect of IL-17RA signaling in the modulation of CD8+ T cell responses based on the fact that CD8+ T cells not only express IL-17RA but also upregulate it in response to cytokines that boost CD8+ T cell responses, such as IL-21 [23,31]. The IL-17RA subunit interacts with IL-17RC to form the functional receptor for both IL-17A and IL-17F (reviewed in [6]), but also with IL-17RD that differentially regulate IL-17A-inducing signaling pathways [32,33]. Therefore, we evaluated by quantitative PCR the amounts of mRNA encoding IL-17RA, IL-17RC and IL-17RD. In agreement with published data, significant amounts of mRNA encoding IL-17RA were detected in spleen CD8+ T cells (Fig 6B). These cells also presented the *Il17rc* and *Il17rd* transcripts but at remarkably lower levels in comparison to *Il17ra*. Studies by flow cytometry confirmed expression of IL-17RA on CD8+ T cells although at different levels according to the CD8+ T cell subset, being maximal in memory cells, and lower in the other subsets. *T. cruzi* infection upregulated IL-17RA expression in naïve and memory CD8+ T cell subsets (Fig 6C).

To directly address the impact of intrinsic IL-17RA signaling in the development of CD8+ T cell responses during *T. cruzi* infection, we performed a series of adoptive transfer experiments. First, we transferred equal number of congenically marked IL-17RA deficient (CD45.2+) and WT (CD45.1+) CD8+ T cells into F1 (CD45.1+/CD45.2+) WT hosts. Recipient mice were infected and at 20 dpi we examined the presence of injected cells in the total and TSKB20-specific CD8+ T cell pools. As expected, no CD45.2+ KO or CD45.1+ WT CD8+ T cells were detected in infected F1 (CD45.1+/CD45.2+) WT hosts that were non-injected (control mice). When analyzing the competitive adoptive transfer experiment, we observed that within the population of injected cells, IL-17RA KO CD8+ T cells were outcompeted by their IL-17RA-sufficient WT counterparts in total and parasite-specific CD8+ T cell subsets all the organs examined including blood and spleen (Fig 6D). To overcome the possible limitations derived from the presence of endogenous WT CD8+ T cells in lymphoreplete hosts, we repeated this competitive experiment in CD8α-/- mice. At 17 dpi, the frequency of IL-17RA KO cells doubled that of WT cells within the total CD8+ T cells (Fig. 6E), likely as result of an increased homeostatic proliferation of polyclonal IL-17RA KO CD8+ T cells that exhibit increased proliferative capacity (Fig 2A). In contrast, WT cells significantly outnumbered IL-17RA KO cells within the TSKB20-specific CD8+ T cell pool, indicating that intrinsic expression of IL-17RA provides a maintenance advantage particularly for parasite-specific CD8+ T cells. Indeed, the ratio between the percentage of WT CD8+ T cells within TSKB20-specific gate and the polyclonal total CD8+ T cells is significantly higher when compared to the same ratio of IL-17RA KO CD8+ T cells (Fig 6F).

Phenotypic evaluation determined IL-17RA KO CD8+ T cells exhibited a higher expression of the inhibitory receptor PD-1 (Fig 6G). Upon antigen specific stimulation with TSKB20, WT CD8+ T cells exhibited a significantly higher polyfunctional effector response in comparison to IL-17RA KO CD8+ T cell from the same host (Fig 6H, left panel and Fig S5). An identical result was observed after polyclonal stimulation (Fig 6H, right panel and Fig S5).

## Discussion

During the last decades, many reports have delineated the immunological mechanisms underlying IL-17-mediated roles in host protection against infection as well as in inflammatory diseases [34,35]. In this paper, we uncover a novel mechanism by which IL-17RA signaling regulates CD8+ T cell responses and promotes protection against a protozoan infection. Our findings provide new elements to elucidate the cytokines and pathways leading to robust CD8+ T cell responses against infections with intracellular microbes and consequently, may have important implications for vaccine and therapy design.

The IL-17 family has a well-established importance in host defense against extracellular bacterial and fungal pathogens and numerous recent studies indicate that it also potentiates immunity to intracellular bacteria, viruses and parasites [3]. So far, the IL-17-dependent protective mechanisms were described to depend on the activation of non-immune and innate immune cells to sustain inflammation [8] and, only indirectly, promote adaptive cellular immune responses [36,37]. In this regard, there are antecedents about the role of IL-17 cytokines in promoting specific CD8+ T cell responses that favored host resistance to Listeria infection and melanoma [38,39]. In these settings, the IL-17-mediated induction of protective CD8+ T cells depended on an enhanced recruitment of cross-presenting dendritic cells but a direct role on CD8+ T cells was not ruled out. Very recently, IL-17 was shown to directly potentiate CD8+ T cell cytotoxicity against West Nile Virus infection [9]. In the same direction, we demonstrate that mice deficient in IL-17RA showed an abortive CD8+ T cell response during *T. cruzi* infection characterized by an early reduction in the number of parasite-specific CD8+ T cells. Concomitantly, infected IL-17RA KO mice showed poor control of the parasite in target tissues such as spleen, liver and heart.

By using different experimental approaches that included a detailed phenotypic, functional and transcriptional profiling of CD8+ T cell arising during *T. cruzi* infection in IL-17RA KO mice as well as *in vitro* and *in vivo* assays, we demonstrated that CD8+ T cell-intrinsic IL-17RA signaling, particularly during the expansion phase, is required to sustain CD8+ T cell survival and therefore, a robust pathogen-specific CD8+ T cell immunity. Still, it remains possible that the cross-talk of IL-17 with other cytokines that regulate CD8+ T cell fate such as IL-21 may be indirectly involved in the effects that we observe. In this regard, Hoft and colleagues described that TCR-transgenic CD4+ T cells specific for an immunodominant peptide of *T. cruzi* and polarized *in vitro* into a Th17 cells potentiated CD8+ T cell immunity in a mechanism dependent on IL-21 but independent on IL-17 [40]. Whether these *in vitro* generated Th17 cells are equivalent those differentiated *in vivo* within the environment of the natural *T. cruzi* infection remains to be established. Indeed, we determined that infected IL-17RA KO mice presented higher concentration of IL-21 in serum than infected WT controls, ruling out deficient IL-21 production as the cause of the altered CD8+ T cell response observed in absence of IL-17RA signaling.

CD8+ T cells from infected IL-17RA KO mice showed increased apoptosis and a transcriptional landscape associated to bio-functions and canonical path of cell death and senescence. In addition, these cells exhibited high expression of multiple inhibitory and death receptors that correlated with reduced *ex vivo* effector responses and a gene signature that positively correlated with those defined for dysfunctional PD-1^high^ CD8+ T cells [41,42]. Altogether these features partially resembled those of dysfunctional CD8+ T cells described in many chronic infections and cancer and characterized by the progressive loss of T cell function, sustained expression of inhibitory receptors and high susceptibility to deletion [43]. Interestingly, we found that the functional CD8+ T cell response observed in *T. cruzi*-infected WT mice (this manuscript and [44]) turned into a poorly-functional response in the absence of IL-17RA signaling. In this regard, although it is conceivable that the increased tissue parasitism in IL-17RA KO mice favored CD8+ T cell exhaustion, infection of WT mice with increasing parasite loads does not induce expression of inhibitory receptors or deletion of parasite-specific CD8+ T cells nor results in lower CD8+ T cell cytotoxic effector function [45]. As cytokines play essential roles in the regulation of exhaustion [20], we performed adoptive transfer experiments that supported the notion that IL-17 signaling directly influenced CD8+ T cell survival and function. In this line, we demonstrated that anti-PD-L1 Abs during the CD8+ T cell expansion phase reinvigorated parasite-specific CD8+ T cell responses in infected IL-17RA KO mice, restoring a robust parasite control and reducing pathology to levels comparable to infected WT controls. Remarkably, we observed that a short regimen of PD-L1 neutralization increased the frequency of parasite-specific CD8+ T cells by two-fold in both treated groups, but the magnitude of the response in infected IL-17RA KO mice was still lower than in WT counterparts. In this regard, it is likely that, given the high levels of several inhibitory receptors in CD8+ T cells elicited by *T. cruzi* infection in absence of IL-17RA, the combination of different anti checkpoint Abs may have stronger effects in these settings, as reported in cancer [46].

So far, the role of checkpoint inhibitors and, particularly the PD-1/PD-L1 axis has been scarcely evaluated during *T. cruzi* infection. One report demonstrated that genetic disruption of this inhibitory pathway increased the effector immune response and consequently, favored parasite control but also promoted cardiac pathology and compromised host resistant to the infection [47]. In this context, our data provide evidences that a PD-L1 checkpoint blockade temporarily limited to the CD8+ T cell expansion phase not only reverses the increased susceptibility to *T. cruzi* absence of IL-17RA signaling but also boosts protective CD8+ T cell immunity during the natural infection. These findings are relevant to a growing area of research aimed at investigating the therapeutic potential of targeting immune checkpoint pathways during chronic infections [48]. In the last years, several reports have shown that immune cell exhaustion or dysfunction is a common finding in human Chagas’ disease, particularly in patients with the most severe conditions [49-52]. There is certain consensus that even the very low parasite load present in tissues may drive, in the long term of the human infection (two decades), the expression of inhibitory receptors and cell dysfunction in T cells, likely resulting in an immune modulatory mechanism and reduced tissue damage in the late stage of infection. Interestingly, many reports have described that patients with moderate and severe chronic chagasic cardiomyopathy present reduced production of IL-17, supporting the notion that this cytokine is protective against cardiac damage [53,54]. Altogether, these and our results support the hypothesis that high production of IL-17 in certain patients may prevent, in the context of persistent parasite levels, the induction of T cell dysfunction and the associated immune dysbalance that cause chronic myocarditis. This hypotheses deserves further evaluation given the relevance for the understanding of the immunopathology during human Chagas’ disease,

From the IPA analysis of the transcriptome of CD8+ T cells arising during *T. cruzi* infection in the absence of IL-17RA, it emerged as potential upstream regulators IRF4 and, at a minor extent, BATF, two TFs reported to be individually required for survival and differentiation of early effector CD8+ T cells [25-27,55-57]. IRF4 promotes CD8+ T cell expansion and differentiation [25], but it also induces the expression of molecules associated with an exhausted status such as Blimp-1, CTLA-4 and IL-10 [58-60]. Furthermore, BATF expression may be triggered by PD-1 ligation and associated with CD8+ T cell exhaustion during chronic viral infection in humans and mice [61]. Very recently, IRF4 and BATF have been shown to form a TCR-responsive transcriptional circuit that establishes and sustains T cell exhaustion during a chronic viral infection [62]. Therefore, it is conceivable that sustained high levels of both TFs could result in the induction of a cell death and exhaustion program as that observed in CD8+ T cells from infected IL-17RA KO mice. How the triangle formed by of loss of IL-17RA signaling, increased expression of BATF and IRF-4 and CD8+ T cell exhaustion is actually linked deserves further studies using genetic dissection approaches.

In conclusion, our results provide evidences for a novel IL-17RA-mediated mechanism that potentiates immunity to an intracellular pathogen like *T. cruzi* by improving adaptive CD8+ T cell responses. This, together with the report showing that IL-17 controls functional competence of NK cells during a fungal infection [63], suggests that cytokines of the IL-17 family would be important for both the innate and the adaptive arms of the cytotoxic response against pathogens. Although, it remains to be determined whether this mechanism is also operative in other infections where IL-17 cytokines have been shown to increase host resistance, these findings have two fundamental repercussions in the field of IL-17-mediated immune responses. The first is that the prolonged targeting of the IL-17 pathways as part of an anti-inflammatory treatment during autoimmune diseases or cancer could potentially lead to severe undesired effects due to defective cytotoxic responses. The second is the notion that promoting the production of IL-17 cytokines during infection or vaccination may help to elicit stronger cytotoxic responses to fight against microbes, and eventually, tumors.

## Material and Methods

### Mice

Mice used for experiments were sex- and age-matched (6 to 10 week-old). C57BL/6 mice were obtained from School of Veterinary, La Plata National University (La Plata, Argentina). IL-17RA KO mice were provided by Amgen Inc. (Master Agreement N° 200716544-002). IL-17A/IL-17F double KO mice [64] were kindly provided by Dr Immo Prinz. CD45.1 C57BL/6 mice (B6.SJL-Ptprca Pepcb/Boy) and CD8α KO mice (B6.129S2-Cd8αtm1Mak/J) were obtained from The Jackson Laboratories (USA). CD45.1 x CD45.2 F1 mice bred in our animal facility. All animals were housed in the Facultad de Ciencias Químicas, Universidad Nacional de Córdoba.

### Animal Welfare

All animal experiments were approved by and conducted in accordance with guidelines of the Institutional Animal Care and Use Committee (IACUC) Facultad de Ciencias Químicas, Universidad Nacional de Córdoba (Approval Number 981/15) (OLAW Assurance number A5802-01). The IACUC adhered to the guidelines from “Guide to the care and use of experimental animals” (Canadian Council on Animal Care, 1993) and “Institutional Animal Care and Use Committee Guidebook” (ARENA/OLAW IACUC Guidebook, National Institutes of Health, 2002).

### Parasites and experimental infection

Bloodstream trypomastigotes of the Tulahuén strain of *T. cruzi* were obtained from BALB/c mice infected 10 days earlier. For experimental infection mice were inoculated intraperitoneally with 0.2 ml PBS containing 5×10^3^ trypomastigotes (usual dose) or doses of 500 and 5×10^4^ trypomastigotes when indicated. For infection of transferred CD8α knockout mice, parasite dose was 1000 trypomastigotes.

### Quantification of parasite DNA in tissues

Genomic DNA was purified from 50 μg of tissue (spleen, liver and heart) using TRIzol Reagent (Life Technologies) following manufacturer's instructions. Satellite DNA from *T. cruzi* (GenBank AY520036) was quantified by real time PCR using specific Custom Taqman Gene Expression Assay (Applied Biosystem) using the primer and probe sequences described by Piron et al [65]. A sample was considered positive for the *T. cruzi* target when CT<45. Abundance of satellite DNA from *T. cruzi* was normalized to GAPDH abundance (Taqman Rodent GAPDH Control Reagent, Applied Biosystem) and expressed as arbitrary units.

### Cells and culture

CD8+ T cells were isolated from the spleen by magnetic negative selection or cell sorting from pools of at least 3-5 mice. For magnetic cell purification the EasySep™ Mouse CD8+ T Cell Isolation Kit (Stemcell Technologies) were used according to the manufacturer’s protocol. For cell sorting, spleen cell suspensions were surface stained and CD3+CD8+ T cells were sorted with a FACSAria II (BD Biosciences). During in vitro studies, CD8+ T cells (2 x 10^5^) purified by magnetic selection were stimulated in 96-well plates coated with anti-CD3/anti-CD28 Abs (eBioscience, 2 μg/ml and 1 μg/ml respectively) and incubated during 24 h in the presence of 100 ng/ml of recombinant IL-17A, IL-17F, IL-17C and IL-17E (ImmunoTools, GmbH) and/or 50ng/mL of IL-21 and TNF (ImmunoTools, GmbH).

### Antibodies and Flow cytometry

Cell suspensions were washed in PBS and incubated with LIVE/DEAD Fixable Cell Dead Stain (eBioscience) during 15 min at RT. Next, cells were washed and incubated with fluorochrome labeled-Abs for 20 min at 4°C (see below for details). To detect *T. cruzi* specific CD8+ T cells, H-2K(b) *T. cruzi* trans-sialidase amino acids 567-574 ANYKFTLV (TSKB20) APC-Labeled Tetramer (NIH Tetramer Core Facility) were incubated 20 min at 4°C before further surface staining with additional Abs. After further surface staining, cells were washed and acquired in a FACSCanto II (BD Biosciences). Blood was directly incubated with the indicated antibodies and erythrocytes were lysed with a 0.87% NH_4_Cl buffer previously to acquisition. For Ki-67 intranuclear staining, cells were first stained on surface, washed and then fixed, permeabilized and stained with Foxp3/Transcription Factor Staining Buffers (eBioscience) following eBioscience One-step protocol: intracellular (nuclear) proteins.

Flow cytometry and/or cell sorting was performed with a combination of the following Abs (BD Biosciences, Biolegend, eBioscience, Life Technologies, Santa Cruz Biotechnology or Cell Signaling): FITC/AF488-labeled anti-mouse: CD8 (53-6.7), Bcl-2 (10C4), IgG (polyclonal goat) and anti-rat IgG (polyclonal goat); PE-labeled anti-mouse: Fas (Jo2), CTLA-4 (UC10-4B9), PD-1 (RMP1-30), Ki-67 (SolA15), IL-17RA (PAJ17R), CD120b (TR75-89), Bim (rabbit. C34C5), rabbit IgG control isotype (DA1E); PECy7-labeled anti-mouse: CD8 (53-6.7), PD-1 (RMP1-30), FAS (Jo2), PerCPCy5.5-labeled anti-mouse: CD3 (145-2C11), CD45.2 (104); PerCP-eFluor 710 labeled anti-mouse: TIGIT (GIGD7); APC-Cy7/AF780-labeled anti-mouse: CD45.1 (A20); Biotin-labeled anti-mouse: BTLA (8F4) and PE-labeled Streptavidin; unlabeled anti-mouse: Bcl-xL (rabbit, 54H7), Bcl-2-associated death promoter (BAD) (rabbit, C-20), Bax (rabbit, P-19).

### Quantification of IL-21

The concentration of IL-21 in the plasma of in plasma of infected WT and IL-17RA KO mice was assessed by ELISA using paired specific Abs (eBiosciences) according to standard protocols.

### Evaluation of Proliferation and Apoptosis

*T. cruzi*-infected WT and IL-17RA KO mice were given BrdU (1 mg/ml, Sigma-Aldrich)/1% sucrose in the drinking water (carefully protected from light) ad libitum. After 5 days, mice were sacrificed and spleen mononuclear cells were collected and stained with surface antibodies. Incorporated BrdU was detected with a BrdU Flow kit according to the manufacturer’s specifications (BD Biosciences).

Apoptosis was determined by Annexin V and 7AAD staining according to the manufacturer’s specifications (BD Biosciences). Mitochondrial depolarization (ψ) was measured by FACS using 50 nM TMRE (Invitrogen) as described [66].

### Determination of CD8+ T cell effector function in vitro

Spleen cell suspensions were cultured during 5 h with medium, 5 μg/ml TSKB20 (ANYKFTLV) peptide (Genscript Inc.) or 50 nM PMA plus 0.5 μg/ml ionomycin (Sigma-Aldrich) in the presence of Monensin (eBioscience) and a PE-labeled anti-CD107a mAb (eBioscience, eBio1D4B). After culture, the cells were surface stained, fixed and permeabilized with BD Cytofix/Cytoperm and Perm/Wash (BD Biosciences) according manufacturer’s instruction. Cells were incubated with FITC-labeled antibody to IFNγ (eBioscience, XMG1.2) and PerCP/APC-labeled antibody to TNF (Biolegend, MP6-XT22) and acquired on FACSCanto II (BD Biosciences).

### Adoptive cell transfer

Competitive adoptive transfer experiments were performed by i.v. injection of CD45.1/CD45.2 F1 recipient WT mice or CD8α knockout mice with a mixture 1:1 of CD8+ T cells purified from spleen of CD45.1+ WT mice and CD45.2+ IL-17RA KO mice. Total 5x10^6^ cells and 15x10^6^ cells were injected in WT and CD8α knockout recipients, respectively. Recipient mice were immediately infected and frequency of the injected CD45.1+ WT and CD45.2+ IL-17RA KO cells within the total T cell population were determined in blood, spleen and liver at different days pi.

### Treatment with neutralizing anti-IL-17A and PD-L1 Abs

To block the PD-1/PD-L1 pathway, infected WT and IL-17RA KO mice were injected i.p. with 200 μg rat anti-mouse PD-L1 antibody (10F.9G2, BioXcell) or total rat IgG isotype control (Jackson Research) every 3 days from 15 dpi until sacrifice at day 21 dpi. Infection matched WT mice were assayed in parallel as controls.

For the in vivo neutralization of IL-17A, infected WT mice were injected i.p. with 200 μg rat anti-mouse IL-17 antibody (17F3, BioXcell) or rat total IgG1 isotype control (MOPC-21, BioXcell) every 3 days from day 13 dpi until sacrifice at 21 dpi. Infection-matched IL-17RA KO mice were assayed in parallel as controls.

### Real-Time quantitative PCR and DNA microarray

RNA from total spleen tissue was purified using TRIzol Reagent (Life Technologies) following manufacturer's instructions. Oligo-dT and a MMLV reverse transcriptase kit (Invitrogen) were used for cDNA synthesis. IL-17RA, IL-17RC and IL-17RD transcripts were quantified by real-time quantitative PCR on an ABI PRISM 7700 Sequence Detector (Perkin-Elmer Applied Biosystems) with Applied Biosystems predesigned TaqMan Gene Expression Assays and reagents according to the manufacturer’s instructions. The following probes were used: Mm00434214_m1 for IL-17RA, Mm00506606_m1 for IL-17RC and Mm00460340_m1 for IL-17RD. For each sample, the mRNA abundance was normalized to the amount of 18S rRNA (Mm03928990_g1) and is expressed as arbitrary units (AU).

For gene-expression analysis, CD8+ T cells were purified by FACS from the spleen of non-infected and 22-day infected WT and IL-17RA KO mice and lysed with TRIzol reagent. Total RNA was extracted with the RNAeasy Mini Kit (Qiagen). RNA quality was verified in an Agilent Bioanalyzer and measured with a Nanodrop 1000 (Thermo Scientific). cDNA was hybridized on Affymetrix Mouse Gene 2.1 ST arrays as described elsewhere.

### Analysis of microarray data

The microarray data from this publication have been deposited to the GEO database (https://www.ncbi.nlm.nih.gov/geo/) and assigned the identifier: GSE104886. Gene expression data was normalized using RMA algorithm on custom Brainarray CDF (v.22.0.0 ENTREZG). Plots of differential expressed genes in spleen CD8 T cells at day 0 post infection and day 22 post infection in WT and IL-17RA KO samples were defined using fold change (|log2FC| >log2(1.2)). Bioinformatic analyses were performed with R software environment. Gene Set Enrichment Analyses (GSEA) between the different spleen CD8 T cell samples were done using specific gene sets from the Molecular Signatures DB (MSigDB). Genes induced by the infection in WT and IL-17RA KO mice but showing significantly higher expression in IL-17RA KO mice were uploaded to Ingenuity Pahtway Analysis (Ingenuity^®^ Systems, www.ingenuity.com) for the analysis of “Disease and bio-function”, “Canonical pathway”, and “Up-stream regulators”. It was considered significantly activated (or inhibited) with an overlap p-value≤0.05 and an IPA activation Z-score as defined under each specific analysis category on the IPA website.

### Statistics

Statistical significance of comparisons of mean values was assessed as indicated by a two-tailed Student’s t test, two way ANOVA followed by Bonferroni’s posttest and Gehan-Breslow-Wilcoxon Test using GraphPad software. P < 0.05 was considered statistically significant.

## Acknowledgements

We thank MP Abadie, MP Crespo, F Navarro, D Lutti, V Blanco, A Romero, L Gatica and G Furlán for their excellent technical assistance. We thank Dr Immo Prinz for providing IL-17A/IL-17F double knockout mice. We acknowledge the NIH Tetramer Core Facility for provision of the APC-labeled TSKB20/Kb and TSKB18/Kb tetramers. We thank A. Rapinat and D. Gentien from Genomic Platform, from the translational research department from Institut Curie, Paris, for the transcriptome experiments.

## Author contributions

JTB designed and performed most of the experiments, analyzed data, and wrote/commented on the manuscript; CLAF, FFV and CR performed experiments, analyzed data and commented on the manuscript; MCR, MCAV, MGS performed experiments and commented on the manuscript; NGN performed microarray experiment; WR analyzed microarray data; EP participated in microarray design and analysis and provided funding; CLM and AG participated in data analysis, commented on the manuscript and provided funding; EVAR supervised the research, designed experiments, wrote the manuscript, and provided funding.

## Supporting Information

**S1 Figure.**
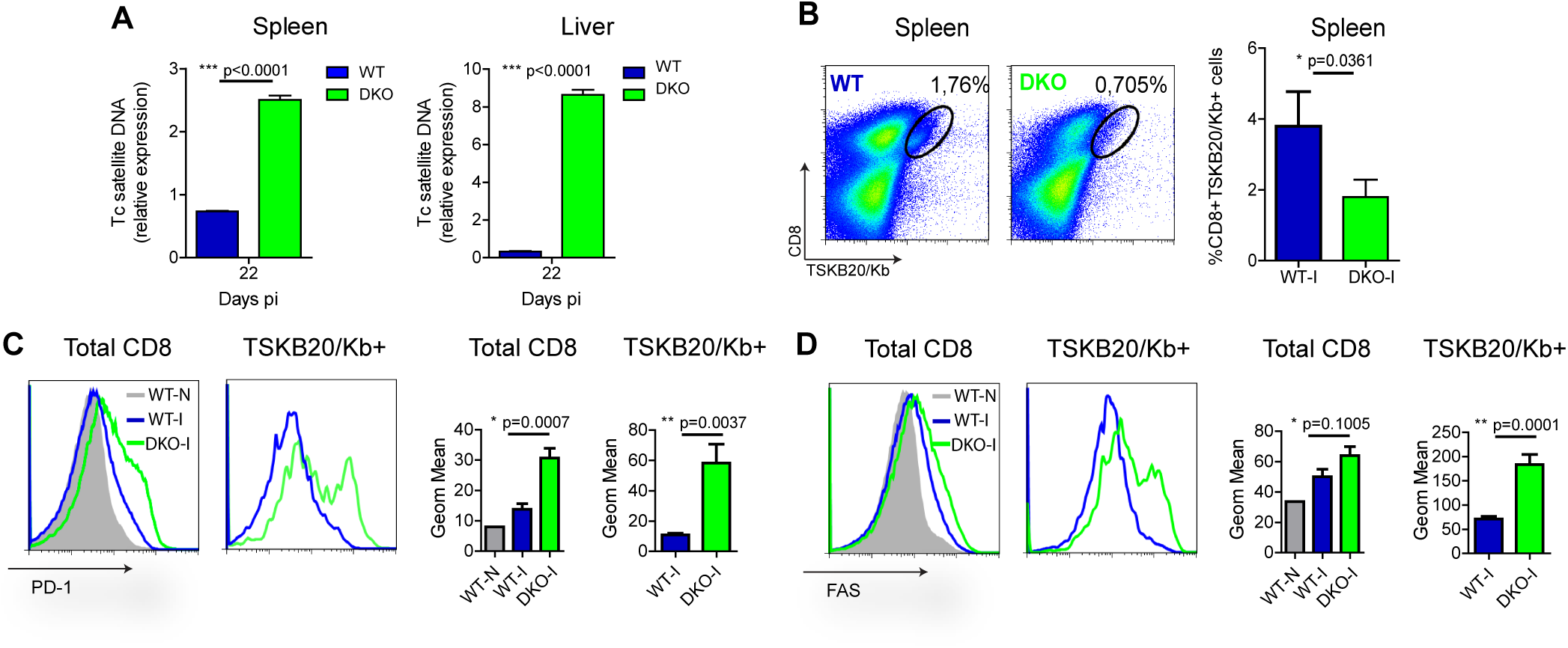
Complementary evaluation of parasite-specific CD8+ T cell responses and tissue parasitism in *T. cruzi*-infected IL-17A/IL-17F DKO mice. **(A)** Relative amount of *T. cruzi* satellite DNA in spleen and liver of infected WT and IL-17A/IL-17F DKO mice determined at 22 dpi. Murine GAPDH was used for normalization. (**B**) Representative plots and statistical analysis of CD8 and TSKB20/Kb staining in spleen of WT and IL-17A/IL-17F DKO mice at 22 dpi. Numbers on plots represent the frequency of TSKB20/Kb+ CD8+ T cells. (**C-D**) Representative histograms and statistical analysis of the geometric mean expression of the inhibitory receptor PD-1 (**C**) and the death receptor CD95/Fas (**D**) in total and TSKB20/Kb+ spleen CD8+ T cells from WT and IL-17A/IL-17F DKO mice at 22 dpi. Grey tinted histogram show staining in CD8+ T cells from non-infected WT mice. Data in statistical analysis (**A-D**) are presented as mean ± SD, N=4–6 mice. P values calculated with two-tailed T test. (**A-D**) Data are representative of at least three independent experiments.

**S2 Figure.**
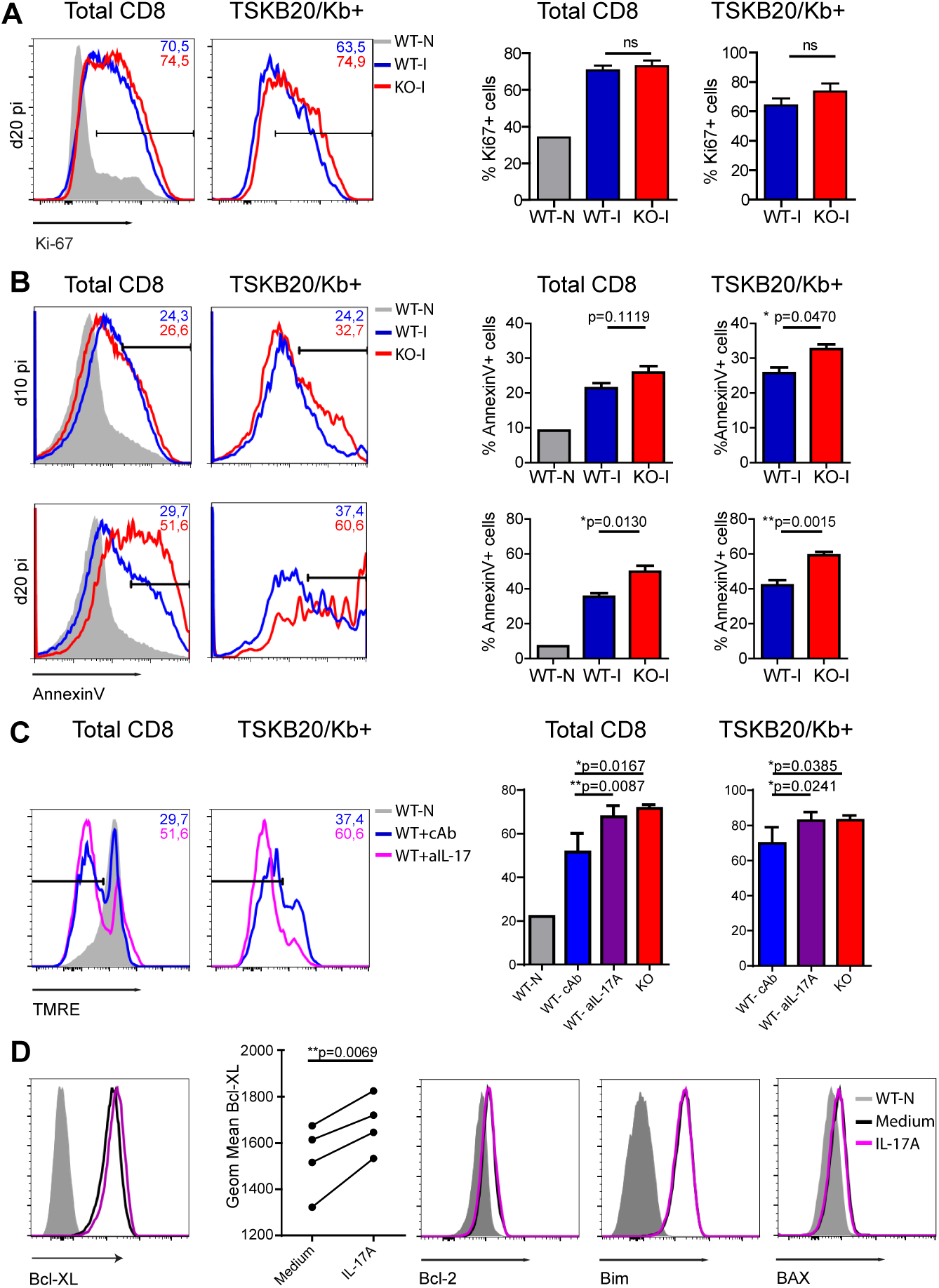
Complementary studies of proliferation and apoptosis in total and parasite-specific CD8+ T cells in WT and IL-17RA KO infected mice. (**A** and **B**) Representative histograms and statistical analysis of Ki-67 (**A**) and Annexin V (**B**) staining within the 7ADD-gate in total (left) and TSKB20/Kb+ (right) CD8+ T cells from the spleen of WT and IL17RA KO mice at 10 dpi (**B**) and/or 20 dpi (**A-B**). Grey tinted histogram show staining in CD8+ T cells from non-infected (N) WT mice. Histograms are representative of one out of five mice. Numbers indicate the frequency of Ki-67+ (**A**) and Annexin V+ (**B**) cells from the corresponding colored group. Bar graphs represent data as mean ± SD, N=4 mice. P values calculated with two-tailed T test. (**C**) Representative histograms of TMRE staining in total (left) and TSKB20/Kb+ (right) CD8+ T cells from the spleen of infected WT mice treated with isotype control or anti-IL-17 as described in Figure 1G. Grey tinted histogram show staining in CD8+ T cells from non-infected WT mice. Numbers indicate the frequency of TMRElow (apoptotic) cells from the corresponding colored group. Histograms are representative of one out of seven mice. Bar graphs in the statistical analysis represent data as mean ± SD, N=7 mice. P values calculated with two-tailed T test. (**D**) Representative histograms of the expression of Bcl-2, Bim and Bax in cultures of purified CD8+ T cells activated during 24h with coated anti-CD3 and anti-CD28 in the presence of medium or IL-17A (100ng/mL) as indicated. (**A-D**) Data are representative of two independent experiments.

**S3 Figure.**
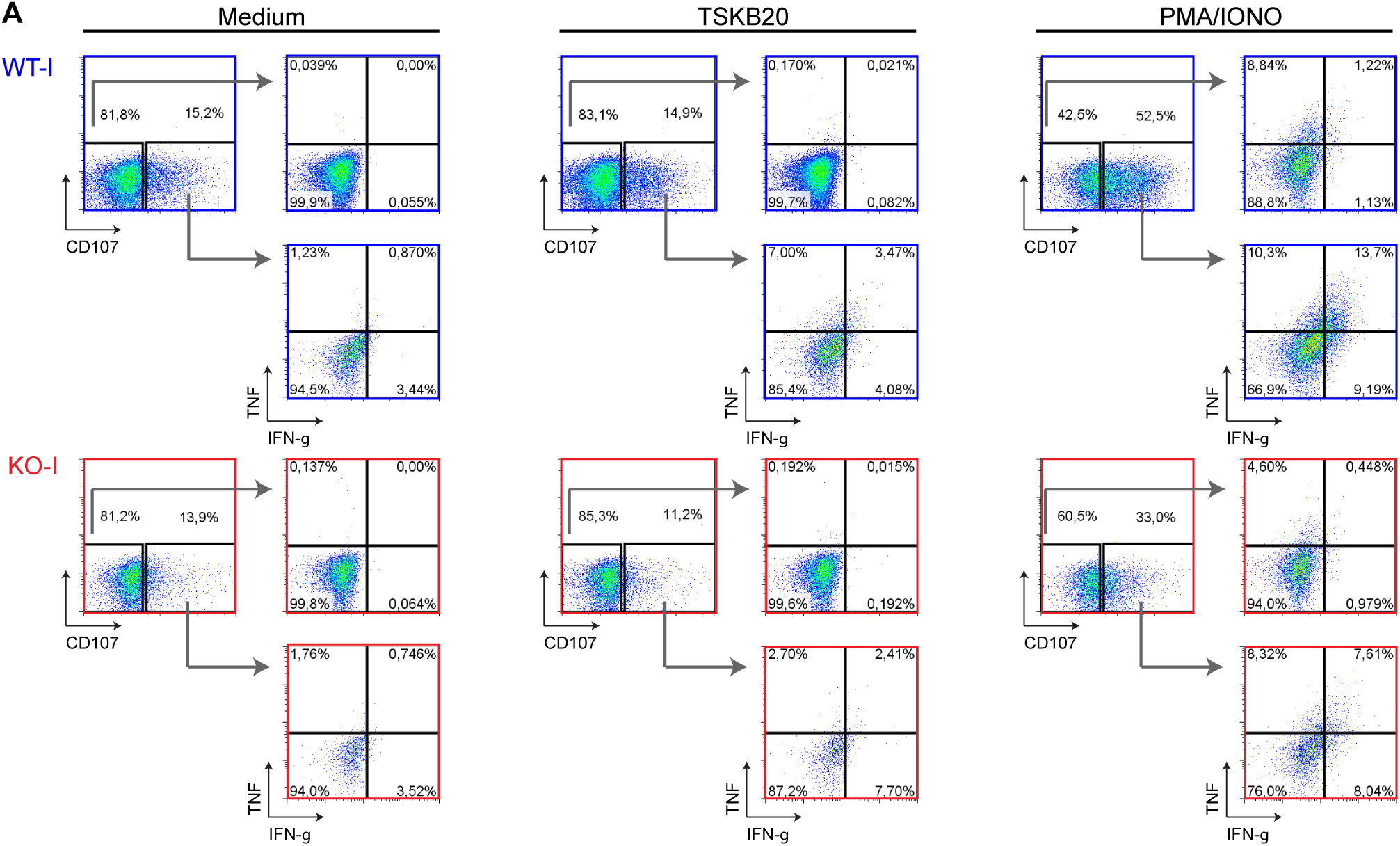
Representative flow cytometry data plots of the evaluation of CD8+ T cell effector function. Representative plots and analysis strategy of the frequency of spleen CD8+ T cells from infected WT and IL-17RA KO mice (22dpi) showing a combination of three and two effector function including expression of CD107a, IFNγ and/or TNF upon 5 h of culture with the indicated stimulation. Plots are representative of one out of five mice. Data are representative of two independent experiments.

**S4 Figure.**
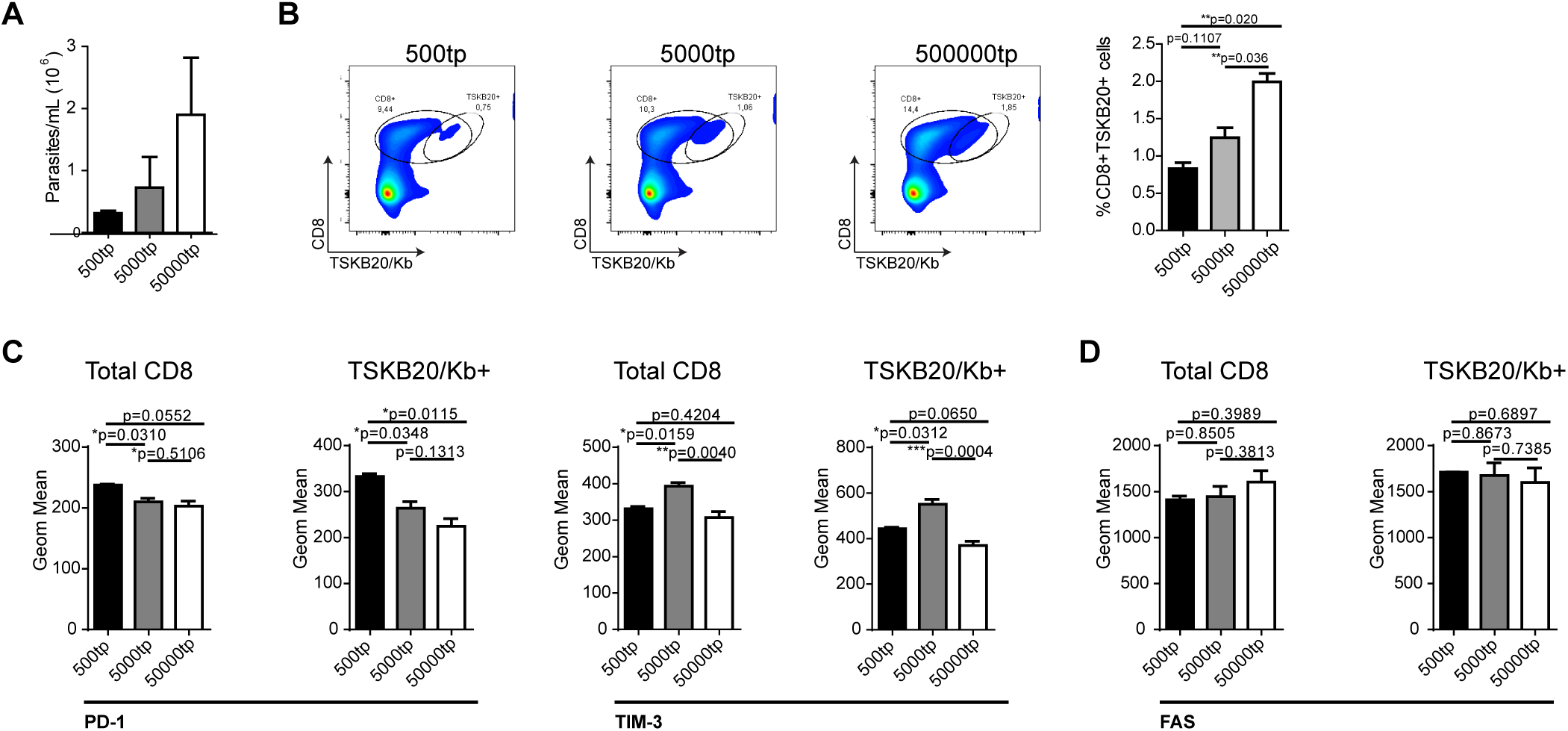
Increasing parasite doses does not diminished the frequency of parasite-specific CD8+ T cells cells or upregulated inhibitory receptors on CD8+ T cells. (**A**) Parasitemia at 22 dpi determined in the blood of WT mice infected with increasing doses of parasites (500, 5000 and 50000 tripomastigotes). (**B**) Representative plots and statistical analysis of CD8 and TSKB20/Kb staining in spleen of WT infected as described in A. (**C** and **D**) Statistical analysis of the geometric mean of expression of inhibitory (**C**) and death (**D**) receptors in total and TSKB20/Kb+ CD8+ T cells from WT mice infected as described in A. Data in statistical analysis are presented as mean ± SD, N=4–6 mice. P values calculated with two-tailed T test. (**A-D**) Data are representative of at least 2 independent experiments.

**S5 Figure.**
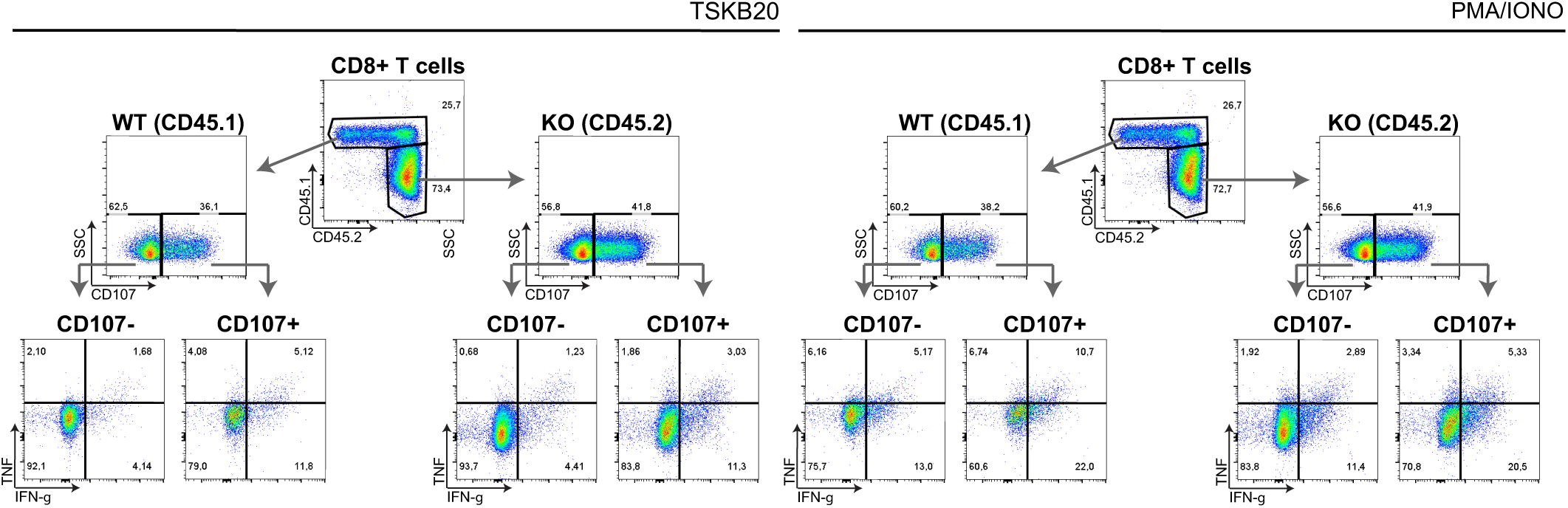
Representative flow cytometry data plots of the evaluation of CD8+ T cell effector function in mice adoptively transferred. Representative plots and analysis strategy of the frequency of CD45.1+ WT and CD45.2+ IL-17RA KO CD8+ T cells from the spleen of CD8α-/- mice adoptively transferred and infected as indicated in figure 7E. The plots show a combination of three and two effector function including expression of CD107a, IFNγ and/or TNF upon 5 h of culture with the indicated stimulation. Plots are representative of one out of four mice. Data are representative of two independent experiments.

